# c-Abl Phosphorylates Plk1 in Facilitating DNA Damage-Induced G2/M Checkpoint Release with a Trade-off of Micronuclei Formation

**DOI:** 10.1101/2025.03.27.645704

**Authors:** Victoria Meltser, Merav Ben-Yehoyada, Julia Adler, Nina Reuven, Yosef Shaul

## Abstract

DNA double-strand breaks (DSBs) pose a critical threat to cellular proliferation and genomic integrity. Upon genotoxic stress, the DNA damage response (DDR) rapidly activates repair pathways and halts cell cycle progression through checkpoint activation. Previously, we demonstrated that DDR-activated c-Abl tyrosine kinase (ABL1) attenuates error-prone late-phase DSB repair. However, the broader functional implications of c-Abl in DDR regulation, and the fate of any residual unrepaired DNA fragments remained poorly understood. Here, we show that c-Abl regulates G2/M checkpoint release by targeting Polo-like kinase1 (Plk1). Depletion or inhibition of c-Abl resulted in increased G2-M accumulation and impaired checkpoint exit. We identified Y217 as a c-Abl phosphorylation site on Plk1, important for Plk1-mediated Claspin destabilization, a key step in G2/M checkpoint release. CRISPR-mediated introduction of the phopsho-silencing Plk1 Y217F mutation or the phospho-mimicking Y217E mutation into cells resulted in impaired and enhanced G2/M checkpoint release, respectively. Intriguingly, c-Abl-mediated G2/M checkpoint release correlated with elevated DNA damage-induced micronuclei (MNi) formation. Depletion or inhibition of c-Abl reduced MNi formation, whereas induction of c-Abl expression increased it, implicating c-Abl as an active effector in both processes. We propose a trade-off model whereby, following the rapid initial repair phase, c-Abl shifts cellular priorities from prolonged, potentially error-prone DSB repair toward cell cycle resumption, thereby promoting G2-M checkpoint exit and DDR deactivation at the cost of increased MNi formation. Our findings describe a novel regulatory DDR axis involving c-Abl and Plk1 and provide mechanistic insights into how DDR termination is orchestrated.

## Introduction

The long-term survival of living organisms depends on their ability to detect and effectively respond to DNA damage (DD). Cellular DNA faces constant threats from both endogenous and exogenous sources (1,2). Among the various types of DNA lesions, double-strand DNA breaks (DSBs) pose the greatest threat to cellular viability (2), as they are highly cytotoxic and can lead to mutations, chromosomal rearrangements, genetic information loss, and genomic instability - a hallmark of cancer (3) (4).

DSBs trigger the immediate activation of the DNA damage response (DDR), a robust and highly coordinated cellular stress response (5), involving the detection of DNA lesions, arrest of cell cycle progression - primarily at the G2/M checkpoint (6) – and initiation of DSB repair. Proper regulation of DDR duration and timely checkpoint deactivation are crucial for maintaining genomic stability and ensuring effective responses to subsequent DNA insults (7,8). While considerable research has focused on DDR activation, including the recruitment of DNA damage sensors, repair factors and checkpoint signaling components (9) significantly less is understood about DDR resolution and its role in determining cell fate (10).

The c-Abl (ABL1) non-receptor tyrosine kinase regulates diverse cellular processes in both the nucleus and cytoplasm (11). Upon genotoxic stress, c-Abl is robustly activated (12) and plays a key role in DNA damage-induced p73-dependent cell death (13). c-Abl also phosphorylates key proteins involved in DNA repair and cell cycle arrest (14,15), and inhibits DNA-PK complex formation with DNA (16). Nevertheless, despite these multiple DDR engagements, the broader scope of c-Abl’s regulatory involvement in the DDR, beyond p73-dependent cell death, remains poorly understood, although its numerous interactions suggest a more extensive role in modulating the cellular response to DNA damage, especially in the context of DSB repair and checkpoint regulation.

DSB repair proceeds in two phases: an initial rapid repair phase, during which most breaks are resolved within minutes to a few hours, followed by a slower, potentially more error-prone repair phase lasting several hours (17). We previously demonstrated that c-Abl downregulates this slower repair phase following ionizing radiation (18). However, the mechanism underlying this downregulation and the fate of the residual unrepaired DSBs remain unclear.

Completion of the second phase of DSB repair correlates with G2/M checkpoint deactivation, particularly following substantial DNA damage (6). This G2/M checkpoint release involves inactivation of the ATR-Chk1, ATM-Chk2 and Wee1 pathways, which mediate DNA damage-induced G2/M arrest (10). Polo-like kinase 1 (Plk1), a key mitotic regulator downstream of Aurora A kinase (AurkA), plays a key role in this process (19–21). Initially, DDR activation inhibits Plk1 (22), suggesting that its reactivation for G2/M checkpoint release depends on late-acting DDR effectors, the identity of which remains unclear.

Here we demonstrate that c-Abl functions as a key effector in this process. Previously we reported that along with c-Abl-mediated attenuation of DSB repair following IR, c-Abl activity correlates with enhanced radiosensitivity and γH2AX foci resolution (18). Since decrease in radiosensitivity and persistence of γH2AX foci have both been associated with prolonged G2/M arrest (23,24), these findings led us to hypothesize that c-Abl activity might be involved in facilitation of G2-M checkpoint release. We now show that c-Abl phosphorylates Plk1 at Y217, potentiating its activity and promoting exit from DNA damage-induced G2/M arrest. Moreover, c-Abl activity was associated with increased micronuclei (MNi) formation, suggesting a potential cellular trade-off to limit error-prone DSB repair. Our findings introduce a novel principle in DNA damage-induced MNi formation, indicating that it serves as a compensatory mechanism balancing genomic integrity with timely DDR deactivation.

## Results

### c-Abl Facilitates Release From the DNA Damage-Induced G2/M Checkpoint

To test whether c-Abl activity affects the duration of G2/M arrest, we inhibited c-Abl pharmacologically, using Imatinib, a well-characterized c-Abl kinase inhibitor, followed by ionizing irradiation (IR) to induce DSBs. Cell cycle distribution was then analyzed via fluorescence-activated cell sorting (FACS) 24 hours post-IR, a time point when cells typically begin to exit DD-induced G2/M arrest. Treatment with the c-Abl inhibitor resulted in an higher proportion of cells in G2/M at 24 hours post-IR compared to untreated controls (Fig. 1A). To determine whether this increase reflected a delay in cell cycle progression, we performed a BrdU pulse-chase assay followed by α-BrdU staining and FACS analysis. In cells with unperturbed c-Abl activity, a large fraction of BrdU-positive cells had transitioned to G1 by 24 hours post-IR, with an even greater proportion reaching G1 by 32 hours post-IR - indicating progression through G2 and mitosis. In contrast, Imatinib-treated cells remained predominantly in G2/M at 24 hours post-IR, with nearly no BrdU-labeled G1 cells. By 32 hours post-IR, a smaller fraction of Imatinib-treated cells had reached G1 relative to controls, strongly suggesting that the c-Abl inhibitor prolonged the G2/M-to-G1 transition (Fig. 1B).

**Figure 1.**
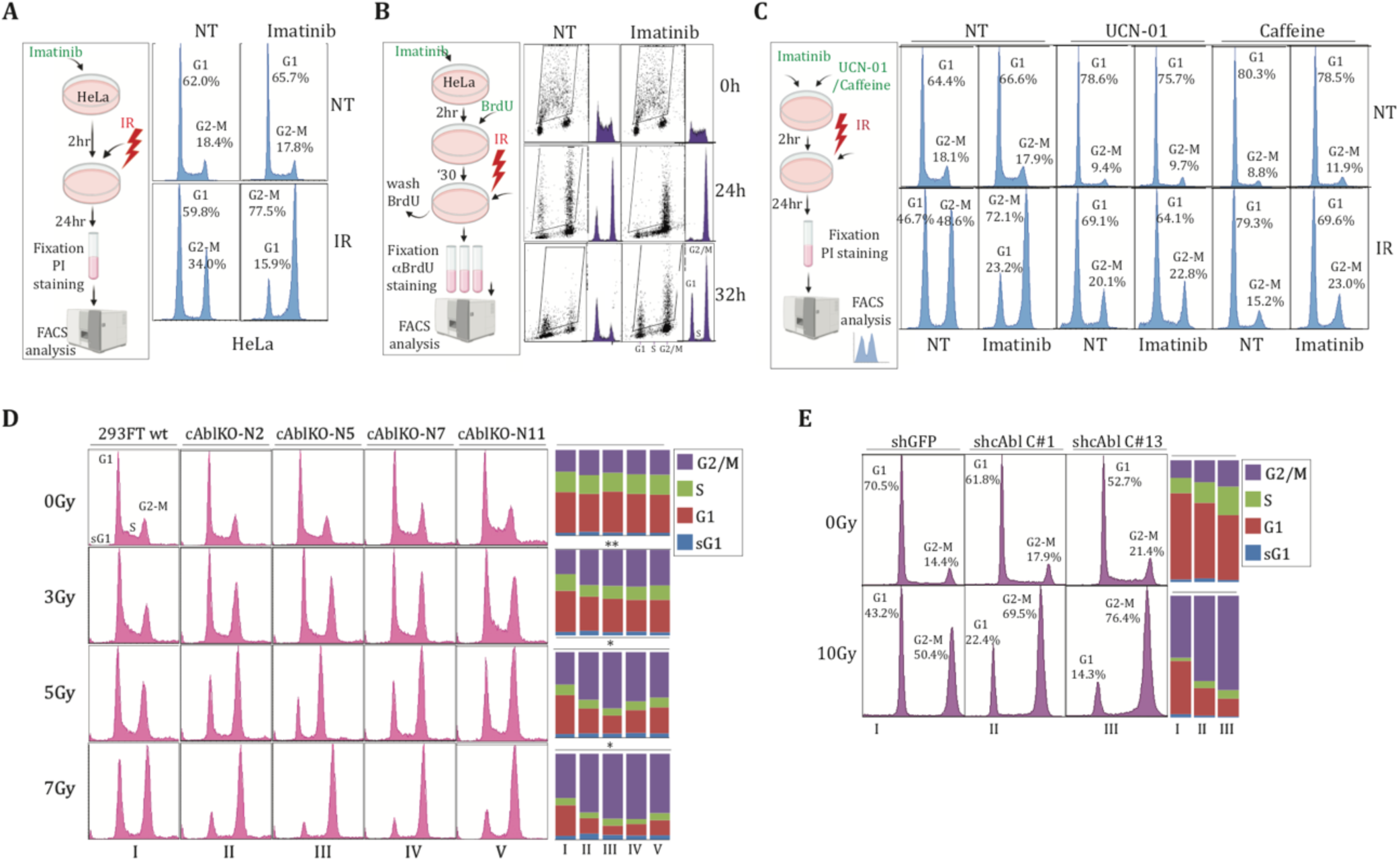
c-Abl tyrosine kinase attenuates DNA damage-induced G2/M arrest. **(A)** HeLa cells were pretreated with Imatinib for 2 hours, irradiated with 10Gy and harvested 24 hours post-IR. Cells were then stained with propidium iodide (PI) and subjected to FACS analysis, as outlined in the experimental scheme (left inset) (**B**) HeLa cells were pretreated with Imatinib for 2hr, pulsed with BrdU for 30 minutes, and irradiated as in (A). Cells were harvested 24 hours post-IR, stained a FITC-conjugated α-BrdU antibody and analyzed by FACS to track the BrdU positive population. (**C**) HeLa cells were left untreated or pretreated with Imatinib for 2 hours, followed by additional treatment with either caffeine or UCN-01 before IR. Cells were harvested 24 hours post-IR, stained with PI and cell cycle distribution was analyzed by FACS. **(D)** Cell cycle profiles of naïve (WT) and c-Abl knockout HEK293FT cells at 24 hours post-IR, exposed to the indicated doses. Statistical significance was determined by one-tailed t-tests comparing the proportion of G2-phase cells in WT 293FT cells and c-Abl KO clones in two independent experiments (**p<0.006, *p<0.015, *p<0.014).

Treatment with the c-Abl inhibitor resulted in an higher proportion of cells in G2/M at 24 hours post-IR compared to untreated controls (Fig. 1A). To determine whether this increase reflected a delay in cell cycle progression, we performed a BrdU pulse-chase assay followed by α-BrdU staining and FACS analysis. In cells with unperturbed c-Abl activity, a large fraction of BrdU-positive cells had transitioned to G1 by 24 hours post-IR, with an even greater proportion reaching G1 by 32 hours post-IR - indicating progression through G2 and mitosis. In contrast, Imatinib-treated cells remained predominantly in G2/M at 24 hours post-IR, with nearly no BrdU-labeled G1 cells. By 32 hours post-IR, a smaller fraction of Imatinib-treated cells had reached G1 relative to controls, strongly suggesting that the c-Abl inhibitor prolonged the G2/M-to-G1 transition (Fig. 1B).

To further validate the effect of c-Abl on G2/M arrest, we overexpressed an active form of c-Abl kinase in cells prior to IR exposure. This resulted in the abrogation of IR-induced G2/M accumulation (Fig. S1A), supporting a link between c-Abl kinase activity and reduced G2/M arrest. Moreover, c-Abl-null mouse embryonic fibroblasts (c-Abl-/- MEFs) were unaffected by Imatinib in terms of IR-induced G2/M accumulation, in contrast to their wild-type counterparts (Fig. S1B), indicating that the observed G2/M delay upon Imatinib pretreatment results specifically from c-Abl inhibition. These findings suggest that c-Abl kinase activity facilitates cell cycle re-entry following DNA damage-induced G2/M arrest, consistent with its role in attenuating late-phase DSB repair.

We next asked whether the G2/M arrest triggered by c-Abl inhibition involves the canonical DD-induced G2/M checkpoint. Specifically, we tested whether Imatinib-induced G2/M accumulation could be reversed by blocking DNA damage checkpoint signaling and preventing activation of the DD-induced G2/M checkpoint. To this end, cells were treated with UCN-01, an inhibitor of Chk1 - a key G2/M checkpoint effector - or with caffeine, which inhibits the major DDR kinases ATM, ATR and DNA-PK, and then exposed to irradiation. Pretreatment with either agent effectively prevented IR-induced G2/M arrest, rendering cells resistant to Imatinib-induced G2/M accumulation (Fig. 1C). These results suggests that the G2/M delay observed under c-Abl inhibition is driven by the canonical DD-induced G2/M checkpoint rather than an alternative pathway.

To further substantiate c-Abl’s role in G2/M arrest following DDR activation, we employed genetic approaches. CRISPR-mediated knockout of c-Abl in HEK293 cells (Fig. S1C) led to pronounced G2/M accumulation following irradiation at various doses, compared to naïve HEK293FT cells (Fig. 1C). Similarly, shRNA-mediated knockdown of c-Abl in U2OS cells (Fig. S1D), also resulted in substantial G2/M accumulation (Fig. 1D). Collectively, these findings highlight c-Abl as a critical regulator of the duration of DNA damage-induced G2/M arrest.

DNA damage-induced arrest in G2/M typically reflects prolonged checkpoint activation, and the duration of this arrest is generally thought to correlate with the timing of G2/M checkpoint release (25). We therefore hypothesized that the extended G2/M arrest observed upon c-Abl depletion or inhibition reflects an active role of c-Abl in driving checkpoint exit. To test this, we used a G2/M checkpoint release assay (Fig. S2A-D). Following cell synchronization, ionizing radiation (IR), and nocodazole treatment, cells were exposed to caffeine - to override continued ATM/ATR checkpoint enforcement - and the extent of G2/M checkpoint release was quantified by measuring the percentage of pH3 (Ser10)-positive mitotic cells.

In c-Abl-knockdown U2OS cells, G2/M checkpoint release was consistently reduced compared to control cells (Fig. 2A). To investigate this further, we utilized doxorubicin, a DNA-damaging agent commonly used in the context of the G2/M checkpoint release assay, and in studies involving c-Abl activation (26). Doxorubicin-induced G2/M arrest was significantly enhanced by Imatinib pretreatment (Fig. S2E), supporting the use of this DNA-damaging agent in studying the role of c-Abl in G2/M checkpoint release.

**Figure 2.**
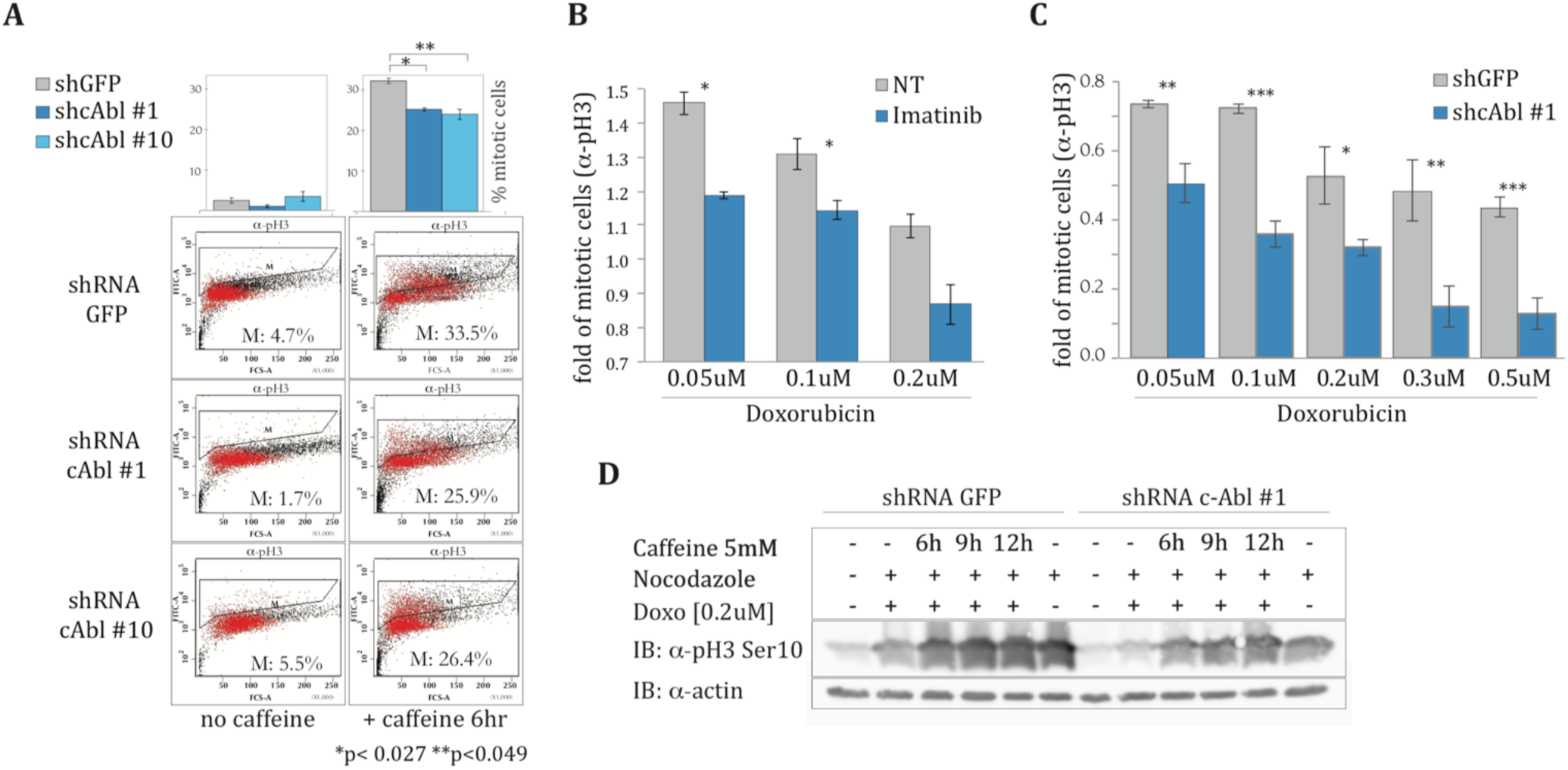
c-Abl tyrosine kinase facilitates G2/M checkpoint release. **(A)** U2OS cells were transduced with control shRNA (shGFP) or two independent shRNAs targeting c-Abl (shcAbl #1, shcAbl #10) were thymidine-synchronized and subjected to a G2/M checkpoint release assay. DNA damage was induced by IR and pH3 (Ser10)-positive mitotic cells were quantified by FACS 24 hours post-IR **(B)** Thymidine-synchronized U2OS cells either pretreated with Imatinib or left untreated and subjected to the G2/M checkpoint release assay. DNA damage was induced with increasing doxorubicin concentrations and the percentage of mitotic cells was determined by FACS **(C)** Thymidine synchronized c-Abl knockdown (shcAbl #1) and control (shGFP) U2OS cells were subjected to the checkpoint release assay. DNA damage was induced with increasing doxorubicin concentrations and the percentage of mitotic cells was assessed by FACS (**p<0.005, ***p<0.0003, *p<0.021, **p<0.006, ***p<0.0004). **(D)** c-Abl knockdown (shcAbl #1) and control (shGFP) U2OS cells were subjected to the checkpoint release assay, and the pH3 (Ser10) mitotic marker was assessed by immunoblot.

Following doxorubicin-induced DNA damage, cells pretreated with Imatinib exhibited significantly reduced G2-M checkpoint release, as evidenced by diminished mitotic entry (Fig. 2B). Similarly, c-Abl knockdown U2OS cells exhibited diminished G2/M checkpoint release when treated with increasing doxorubicin concentrations, compared to control knockdown cells (Fig. 2C). Notably, these knockdown cells also displayed a more pronounced doxorubicin-induced G2/M arrest (Fig. S2F). Immunoblot analysis further confirmed decreased phospho-histone pH3 (Ser10) levels in c-Abl-knockdown cells subjected to the checkpoint release assay (Fig. 2D). Collectively, these findings indicate that c-Abl tyrosine kinase actively promotes G2/M checkpoint release, thereby limiting the duration of DD-induced G2/M arrest.

### c-Abl Phosphorylates Polo-like Kinase 1 (Plk1) at Y217, Potentiating G2/M Checkpoint Release

Plk1 is a key regulator of the transition from DNA damage-induced G2/M arrest to G2/M checkpoint release and subsequent cell cycle re-entry (7). Plk1 activation during G2/M checkpoint release requires phosphorylation at T210 within the Plk1 transactivation loop (T-loop)) by Aurora A kinase (20). Seven residues downstream of T210 lies a conserved tyrosine (Y217), which resides within a sequence matching a c-Abl phosphorylation consensus motif (Y-x-x-P) (27). Sequence alignment of Plk1 across species confirms the high conservation of this region, encompassing both T210 and Y217 (Fig. 3A), underscoring the potential functional significance of Y217. Post-translational modification databases (28) report significant Y217 phosphorylation in multiple cancer cell lines, particularly in CML-derived K562 cells, which exhibit elevated c-Abl kinase activity due to presence of the BCR-ABL fusion gene. These observations led us to hypothesize that c-Abl phosphorylates Plk1 at Y217, thereby regulating Plk1 function in the DNA damage response.

**Figure 3.**
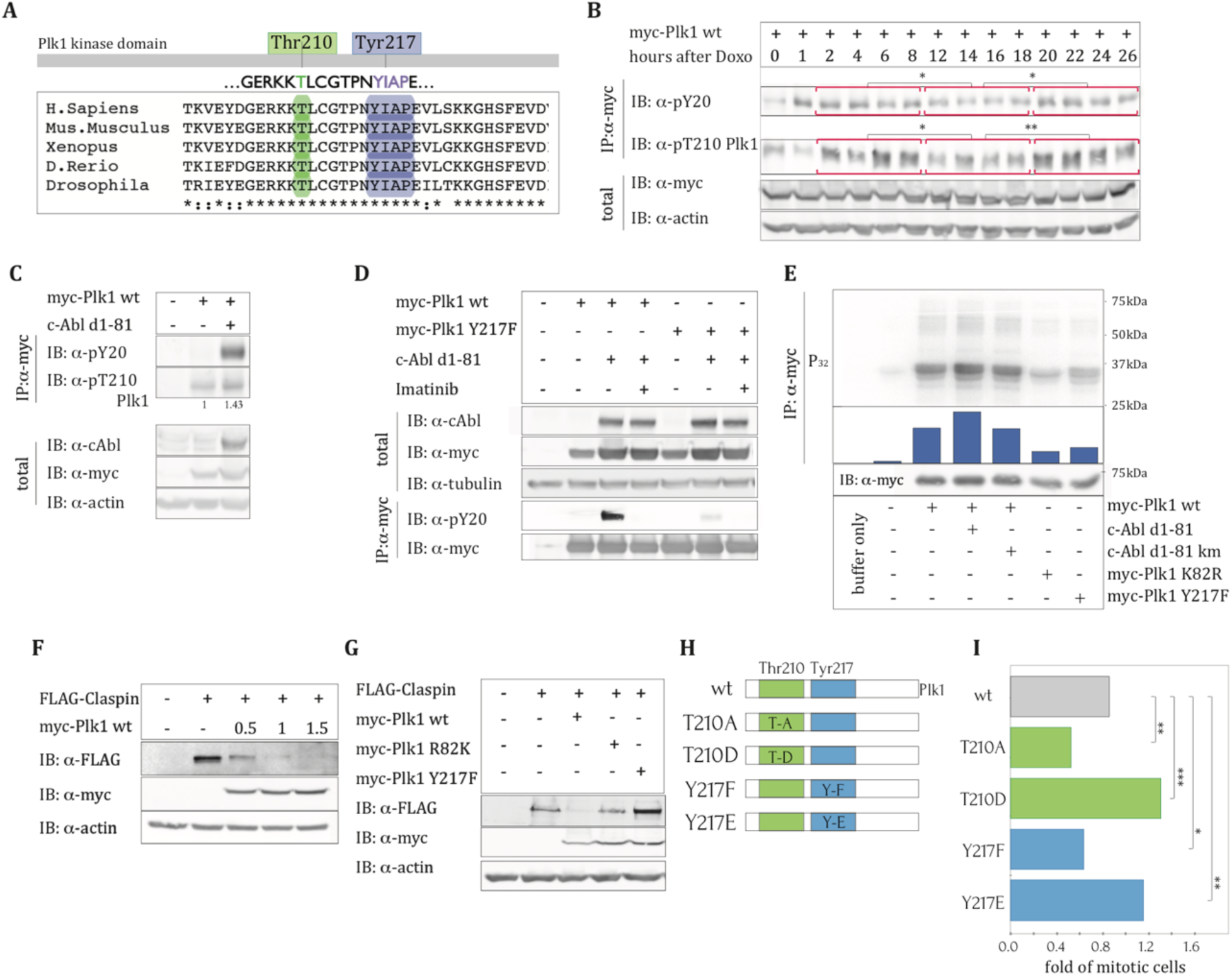
c-Abl Tyrosine Kinase Phosphorylates Plk1 on Tyr217, Critical for Plk1 Kinase Activity and G2/M checkpoint release. **(A)** Sequence alignment of the Plk1 gene sequence across species, highlighting the high conservation of the Plk1 kinase domain and the Plk1 T-loop, including T210 and Y217. Y217 resides within a c-Abl phosphorylation consensus motif **(B)** Time course of Plk1 activation following doxorubicin-induced DNA damage. HEK293 cells were transfected with myc-Plk1 and treated with doxorubicin for 1 hour, 24 hours post-transfection. Cells were harvested at the indicated time points, and lysates were analyzed by immunoblotting with α-myc, α-phosphotyrosine (PY20) to detect Plk1 tyrosine phosphorylation and α-Plk1-T210, to assess Plk1 activation. Time points were grouped into three intervals (2-8 hours, 12-18 hours, and 20-26 hours), and average intensities were quantified. Statistical significance was determined by a one-tailed t-test (*p<0.024, *p<0.012, *p<0.04, **p<0.005). **(C)** HEK293 cells were co-transfected with myc-Plk1 WT and either an empty vector or constitutively active c-Abl (c-Abl Δ1-81). At 24 hours post-transfection, Plk1 was immunoprecipitated with α-myc, followed by immunoblotting with PY20 and α-pT210, to detect Plk1 tyrosine phosphorylation and activation, respectively **(E)** HEK293 cells were transfected with myc-Plk1 WT or myc-Plk1 Y217F, with or without co-expression of c-Abl Δ1-81, to assess the importance of Y217 in c-Abl-mediated Plk1 phosphorylation. Plk1 was immunoprecipitated with α-myc, followed by immunoblotting with α-pY20. Imatinib was added to the indicated samples 2 hours prior to harvesting. **(F)** In-vitro kinase assay. HEK293 cells were transfected with the indicated constructs, and Plk1 was immunoprecipitated (α-myc). Immunoprecipitates were subjected to an in-vitro kinase assay followed by immunoblotting, as described in the Methods section. **(G)** HEK293 cells were transfected with increasing amounts of myc-Plk1 WT and FLAG-Claspin. Lysates were immunoblotted with the indicated antibodies to validate the effect of Plk1 expression on Claspin levels. **(H)** HEK293 cells were transfected with the indicated myc-Plk1 constructs, along with FLAG-Claspin to assess the impact of the phospho-silencing Plk1 Y217F mutation on Claspin levels. **(I)** Schematic representation of U2OS clones generated by CRISPR-mediated genome editing, carrying the specified Plk1 mutations **(J)** U2OS clones carrying these Plk1 mutations were treated with doxorubicin for 1 hour to induce DNA damage, then subjected to the checkpoint release assay detailed in Fig.2. Cells were stained for phospho-histone H3 (pH3 Ser10), and analyzed by flow cytometry. Data were normalized to untreated controls. Results are presented as mean ± SEM from three independent experiments. Statistical significance was evaluated using a one-tailed t-test (**p<0.016, ***p<0.0075, *p<0.0058, **p<0.0008).

To examine the kinetics of Plk1 tyrosine phosphorylation following doxorubicin-induced DD and its potential correlation with Plk1 T210 phosphorylation, we conducted a time-course experiment. Cells were transfected with myc-tagged Plk1 (myc-Plk1 WT), treated with doxorubicin for 1 hour, and harvested at 2- to 4-hour intervals, over a 24-hour period. Lysates were to immunoprecipitated with an α-myc and analyzed by immunoblotting with α-pT210-Plk1 (to assess Plk1 activation) and the phopshotyrosine-specific antibody α-pY20 (to detect Plk1 tyrosine phosphorylation). Plk1 tyrosine phosphorylation exhibited a “wave-like” pattern that closely mirrored fluctuations in T210 phosphorylation, peaking at approximately 20 hours post-DD induction - coinciding with the timing of G2/M checkpoint release and mitotic entry (Fig. 3B). A comparable pattern of Pk1 tyrosine phosphorylation was observed following IR-induced DNA damage, within a comparable timeframe (Fig. S3A). These observations hint toward a possibility that Plk1 tyrosine phosphorylation regulates Plk1 function during G2/M checkpoint release.

Next, we examined whether c-Abl affects both tyrosine and threonine phosphorylation of Plk1. Cells were co-transfected with myc-Plk1 WT and a constitutively active form of c-Abl (c-Abl Δ1-81), followed by immunoprecipitation (α-myc) and immunoblotting (Fig.3C). In the presence of active c-Abl, Plk1 displayed strong tyrosine phosphorylation (pY20) alongside elevated T210 phosphorylation (Fig. 3C), suggesting that c-Abl may not only phosphorylate Plk1 but also promote Plk1 T210 phosphorylation, potentially enhancing Plk1 activation under conditions where c-Abl is active.

To determine whether c-Abl directly phosphorylates Plk1 at Y217, constitutively active c-Abl (c-Abl Δ1-81) was co-expressed with either myc-Plk1 WT or the Y217F mutant (myc-Plk1 Y217F) (Fig. 3D). Immunoprecipitation of myc-Plk1 followed by immunoblotting with a general phosphotyrosine antibody (pY20) revealed robust tyrosine phosphorylation of Plk1 when co-expressed with active c-Abl, but this was nearly abolished in the Y217F mutant (Fig. 3D), indicating that Y217 is the primary Plk1 tyrosine residue targeted by c-Abl. Furthermore, treatment with Imatinib effectively blocked Plk1 tyrosine phosphorylation, supporting that this modification is mediated by c-Abl kinase activity. Collectively, these results demonstrate that c-Abl directly phosphorylates Plk1 at Y217, and that pharmacological inhibition of c-Abl prevents Plk1 tyrosine phosphorylation, further supporting a direct regulatory link between c-Abl and Plk1.

To evaluate whether c-Abl–mediated tyrosine phosphorylation directly affects Plk1’s catalytic function, we performed an in-vitro kinase assay (Fig. 3E). The kinase activity of immunoprecipitated Plk1 was measured using ^32^P radiolabeling, with Casein kinase (CSK) - a generic substrate commonly used to assess Plk1 activity (29), in the presence or absence of active c-Abl. While Plk1 WT alone displayed substantial kinase activity, this activity was markedly enhanced by co-expression with active c-Abl (c-Abl Δ1-81), but not with the catalytically inactive c-Abl kinase mutant (c-Abl Δ1-81 km). In contrast, both the kinase-dead Plk1 R82K mutant and the phospho-silencing Plk1 Y217F mutant exhibited significantly reduced baseline activity compared with Plk1 WT (Fig. 3E). These findings suggest that c-Abl mediated phosphorylation positively modulates Plk1’s catalytic function and highlight the importance of Plk1 Y217 phosphorylation for Plk1 kinase activity.

To evaluate the functional significance of Plk1 Y217 phosphorylation in G2/M checkpoint release, we examined the effect of Plk1 on Claspin, a key regulator of G2/M checkpoint maintenance. Claspin is essential for ATR-mediated Chk1 activation during both the initiation and maintenance of the DD-induced G2/M checkpoint (30). During G2/M checkpoint release, Plk1-mediated phosphorylation triggers Claspin degradation via the SCF^β-TrCP^ ubiquitin ligase, (31–33). Co-expression of increasing amounts of Plk1 WT with FLAG-Claspin resulted in a dose-dependent reduction in Claspin levels (Fig. 3F). Importantly, whereas Plk1 WT eliminated Claspin expression, the kinase dead Plk1 K82R had little effect on Claspin levels and the phospho-silencing Plk1 Y217F not only failed to reduce Claspin but instead elevated it above baseline levels (Fig. 3G), strongly suggesting that Plk1 Y217 phosphorylation is functionally critical for Plk1-mediated Claspin destabilization and, by extension, for efficient G2/M checkpoint release.

Finally, we used CRISPR-mediated genome editing to introduce phospho-silencing (Y217F or T210A) or phospho-mimicking (Y217E or T210D) Plk1 mutations into U2OS cells (Fig. 3H). The cells were then assessed for their ability to undergo G2/M checkpoint release, using the checkpoint release assay. As expected, the T210A mutation impaired G2/M checkpoint release, while T210D enhanced it (Fig. 3I), Notably, the Y217F mutation led to reduced G2/M checkpoint release, like T210A, while the Y217E mutation mirrored the T210D mutation, promoting checkpoint exit (Fig. 3I). Together, these findings highlight Plk1 Y217 phosphorylation as crucial for G2/M checkpoint release, with c-Abl as a key modulator of Plk1 activity during late DDR.

### c-Abl Tyrosine Kinase Facilitates DNA Damage-Induced Micronuclei Formation

When cycling cells are exposed to DNA damage in the form of DSBs, they frequently form micronuclei (MNi) (34). Notably, disruption or premature override of the G2/M DNA damage checkpoint leads to a marked increase in MNi formation (35), suggesting that MNi formation reflects a balance between DSB repair efficiency and the duration of DNA damage-induced G2/M arrest. We hypothesized that c-Abl-mediated facilitation of G2/M checkpoint release, as demonstrated in this study, along with its role in moderating DSB repair (18), may lead to incomplete DSB repair and residual unrepaired DSBs, thereby promoting DD-induced MNi formation.

To investigate this possibility, U2OS cells were pretreated with Imatinib prior to irradiation, and MNi formation was quantified 48 hours post-IR. Imatinib-treated cells exhibited a significantly lower percentage of MNi compared to untreated controls (Fig. 4A and Fig. S4A). Similar effects were observed in MCF10A cells (Fig. 4B and Fig. S4B). We next examined DD-induced MNi formation in cells expressing either wild-type c-Abl kinase (c-Abl WT) or a kinase-inactive c-Abl mutant (c-Abl km). Induction of c-Abl WT expression resulted in a significant increase in MNi formation compared to both uninduced cells and those induced to express c-Abl km (Fig. 4C and Fig. S4C).

**Figure 4.**
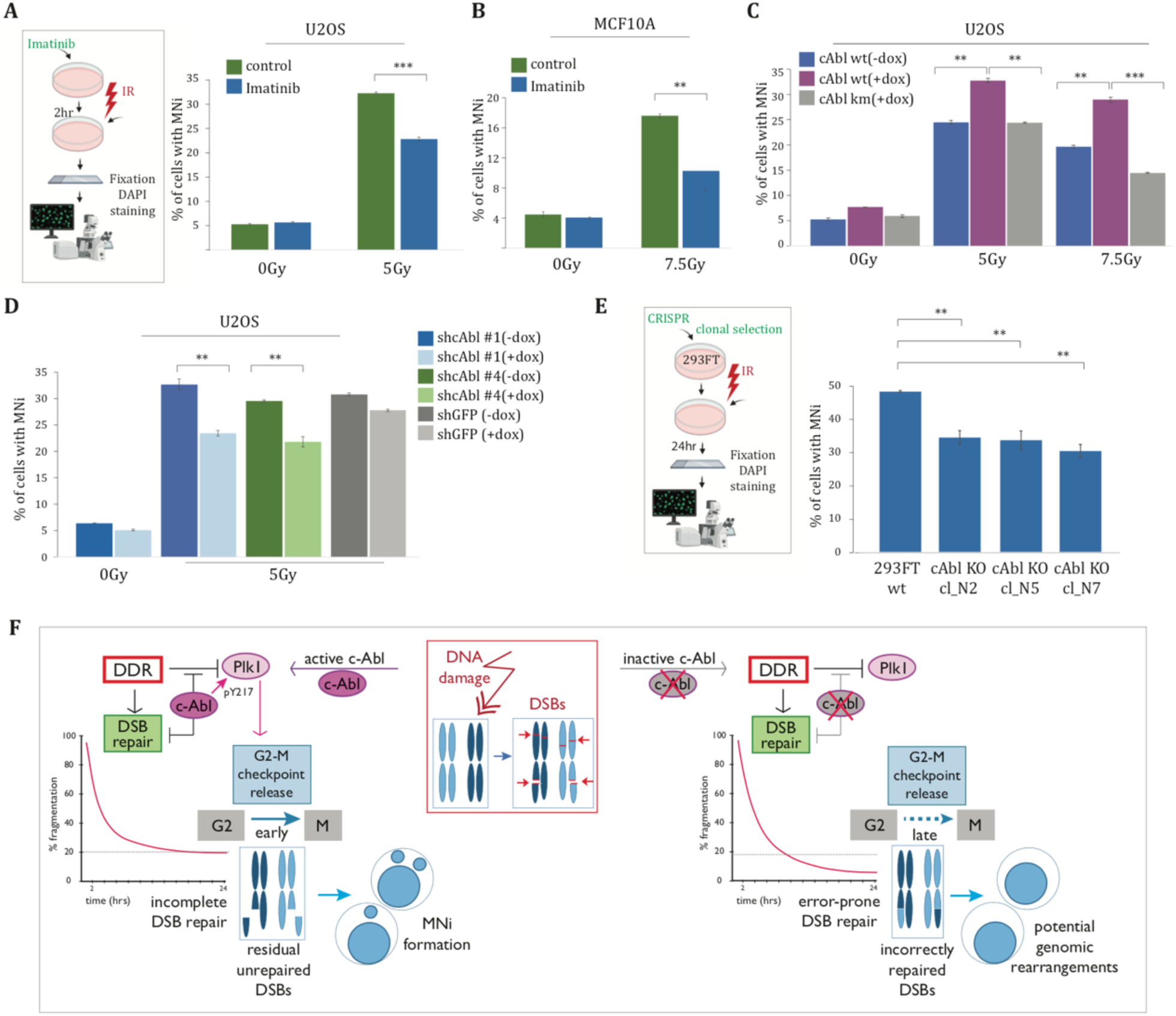
c-Abl tyrosine kinase promotes micronuclei formation following DNA damage. **(A)** U2OS cells were pretreated with Imatinib and DNA damage was induced by IR (10Gy), followed by fixation and DAPI staining 48 hours post-IR. Images were captured using a fluorescent microscope and micronuclei quantification was performed as described in the Methods section. **(B)** MCF10A cells were treated as in (A), collected 72 hours post-IR for DAPI staining and micronuclei quantification. **(C)** U2OS cells expressing a doxycycline-inducible wild-type c-Abl (c-Abl WT) or c-Abl kinase-deficient mutant (c-Abl km) were treated with doxycycline and exposed to the indicated IR doses. Cells were collected 48hr post-IR, for DAPI-staining and micronuclei quantification. Statistical significance was determined using a one-tailed t-test (**p<0.0021, **p<0.0012, **p<0.0018, ***p<0.0007). **(D)** U2OS cells expressing doxycycline-inducible shRNA targeting c-Abl or GFP (control), were left untreated or induced with doxycycline, followed by irradiation at 5Gy. Cells were harvested 48hr post-IR, stained with DAPI and analyzed for micronuclei. Statistical significance was determined using a one-tailed t-test (*p<0.0074, **p<0.0083) **(E)** Control HEK293FT cells and c-Abl knockout clones were irradiated at 10Gy and collected 24 hours post-IR. Micronuclei were quantified as described in the Methods section. Statistical significance was determined using a one-tailed t-test (**p<0.0054, **p<0.0092, **p<0.0032). **(F)** The model illustrates the role of c-Abl in the DNA damage response (DDR), focusing on the involvement of c-Abl in DSB repair and of the c-Abl-Plk1 axis in regulating and G2/M checkpoint release, with consequences for cellular fate. This model suggests that c-Abl acts as a switch between extended DSB repair (potentially introducing repair errors and genomic rearrangements) and G2/M checkpoint release (favoring cell division at the cost of MNi).

Lastly, MNi formation was assessed following c-Abl depletion. Inducible knockdown of c-Abl using two distinct shRNAs in U2OS cells led to a significant reduction in MNi formation (Fig. 4D and Fig. S4D). Additionally, MNi formation was examined in three CRISPR-generated c-Abl knockout clones. While both wild-type and knockout cells formed MNi after DNA damage, c-Abl knockout cells consistently exhibited fewer MNi compared to their wild-type counterparts (Fig. 4E and Fig. S4E). Collectively, these findings suggest that c-Abl mediated facilitation of G2/M checkpoint release following DSBs represents a functional trade-off: while promoting the cell cycle resumption, c-Abl may also contribute to the persistence of residual unrepaired DNA fragments, ultimately leading to increased MNi formation.

## Discussion

A central question in the DNA damage field is how cells regulate the release from the DD-induced G2/M checkpoint. DSBs are considered the most genotoxic type of DNA lesions and maintaining genome integrity in response to such damage requires robust, coordinated cellular mechanisms. The dual-phase nature of DSB repair (17), comprising an initial rapid rejoining of DNA ends followed by a more extended, potentially error-prone, repair phase for complex lesions, underscores the necessity for finely tuned regulatory mechanisms. Concurrent activation of the DD-induced G2/M checkpoint upon exposure to DNA damage ensures that cell cycle progression is halted to provide sufficient time for DSB repair. However, the mechanisms responsible for DDR resolution, which comprises timely checkpoint release and resumption of cell cycle progression, remain incompletely understood. Given that Polo-like kinase 1 (Plk1) plays a central role in orchestrating G2/M checkpoint release, elucidating the DDR mechanisms that restore Plk1 activity following its initial inhibition in response to DD is of critical importance.

Our findings suggest that c-Abl tyrosine kinase (ABL1), which is activated by key DDR regulators under genotoxic stress (12,16,36,37), functions as a regulatory switch to promote G2/M checkpoint release. We demonstrate that c-Abl facilitates checkpoint exit through its interaction with Plk1, thereby coordinating the balance between continued DSB repair and the inactivation of DDR signaling. Inhibition or depletion of c-Abl results in a pronounced and sustained G2/M arrest and in reduced G2/M checkpoint release. This effect can be counteracted by abrogating G2-M checkpoint activation, via inhibition of the checkpoint kinase Chk1 or of the key DDR kinases responsible for c-Abl activation in the DDR (12,37). These findings support a model in which c-Abl plays a pivotal role in regulating the exit from the canonical DD-induced G2/M checkpoint.

Our study identifies a previously uncharacterized c-Abl phosphorylation site on Plk1, Y217, a highly conserved residue positioned near the activating T210 residue within the Plk1 activation loop. While c-Abl has been reported to phosphorylate Plk1 at Y245 in the context of cervical cancer (38), our findings establish Y217 as the primary c-Abl target site on Plk1, critical for Plk1 function in the DDR. The physiological relevance of Y217 as a c-Abl target is further supported by reports of elevated Plk1 Y217 phosphorylation in multiple chronic myeloid leukemia (CML) samples (28), which exhibit elevated c-Abl tyrosine kinase activity due to the presence of the BCR-ABL fusion oncoprotein (39,40).

Kinetic analyses indicate that following DDR activation Plk1 Y217 phosphorylation temporally correlates with T210 phosphorylation, reinforcing a model whereby c-Abl-mediated phosphorylation positively regulates Plk1 activity to drive G2-M checkpoint release. In line with this, we find that constitutively active c-Abl bolsters Plk1 kinase activity in-vitro, and the phospho-mimetic Y217E mutant augments G2-M checkpoint release, while the phospho-silencing Y217F mutation prevents Plk1-driven Claspin destabilization and attenuates G2/M checkpoint release, all of which underscore the functional significance of Y217 in Plk1-mediated G2/M checkpoint exit. We suggest that c-Abl acts as a late DDR off-switch, which regulates DDR deactivation by targeting Plk1 at Y217, promoting its activation and facilitating G2/M checkpoint release.

Our study also reveals a connection between c-Abl-mediated G2/M checkpoint release and increased micronuclei (MNi) formation. DD-induced MNi typically arise from acentric chromosomal fragments that fail to reintegrate into the nucleus during cell division (34), a process influenced by the duration of G2/M arrest (41). While often associated with genomic instability, MNi may also function as a protective mechanism to sequester irreparably damaged chromosomal fragments, thereby limiting potentially error-prone repair. The physiological importance of MNi has garnered increasing attention (42–47), underscoring the importance of understanding the mechanisms that regulate their formation. Moreover, recent evidence indicates that MNi activate the cGAS–STING pathway, a cytosolic DNA-sensing mechanism integral to the innate immune response (43,44) with significant implications for tumor immunosurveillance and inflammatory diseases. Understanding how c-Abl–dependent G2/M checkpoint release influences MNi formation—and, by extension, potential immune activation through cGAS–STING—may therefore offer important insights into cancer progression and therapeutic interventions.

Our findings highlight two potential cell fate decisions that emerge following DNA damage, with the duration of G2/M arrest at the center of this pivotal decision-making point (Fig. 4F). Placed in the broader context of DDR regulation, c-Abl’s ability to expedite G2/M checkpoint release - coupled with its previously described attenuation of late-phase DSB repair (14,18), reveals a nuanced trade-off. On one hand, c-Abl limits prolonged and potentially more error-prone DSB repair while also limiting extended cell cycle arrest, promoting the resumption of cell cycle progression, and averting outcomes associated with prolonged arrest (mutations resulting from error-prone DSB repair, senescence, apoptosis etc). On the other, repair attenuation and facilitation of checkpoint exit may result in residual unrepaired DNA fragments, contributing to MNi formation and elevating the risk of chromosomal and genomic instability. While additional quality control mechanisms may mitigate some of these outcomes, it seems that c-Abl-mediated G2/M checkpoint release reflects a strategic trade-off between maintenance of cell viability through timely cell cycle re-entry and the risks associated with incomplete DSB repair.

## Materials and Methods

### Cell Lines & Reagents

Human osteosarcoma U-2 OS cells (referred to as U2OS), human embryonic kidney HEK-293 cells and human cervical carcinoma HeLa cells and mouse embryonic fibroblasts (MEFs) (WT and c-Abl null) were maintained in Dulbecco’s modified Eagle’s medium supplemented with 10% heat-inactivated fetal calf serum (FCS), 50 μg/ml penicillin, and 50 μg/ml streptomycin. Cells were cultured at 37 °C in a humidified atmosphere and in a 5% CO2 environment. To induce DNA damage, where indicated, cells were exposed to irradiation using an X-RAD320 irradiator, or to Doxorubicin (0.2 micromolar). Imatinib (STI-571) (Novartis, Switzerland) was added at a concentration of 10mM 2hr prior to irradiation. Mouse monoclonal α-phospho-H2AX antibody (Upstate-Millipore, clone JBW301) was used at a concentration of 1:900 for immunofluorescence and 1:10,000 for immunoblotting. Other antibodies used were - α-c-Abl (K-12), α-phospho-Tyrosine (PY20), α-β-tubulin, α-Wee1 - all from Santa Cruz Biotechnology (Santa Cruz, CA, USA), α-phospho-Histone3-Ser10 (Upstate-Millipore), α-phospho-Chk1-Ser345 (Cell Signaling), α-phospho-Plk1-Thr210 (Abcam), α-FLAG (Sigma-aldrich), α-myc mouse monoclonal antibodies were generated by the Antibody Laboratory at the Weizmann Institute, Horseradish peroxidase-conjugated secondary antibodies were from Jackson ImmunoResearch Laboratories (West Grove, PA, USA)

### Flow Cytometry

For cell cycle analysis, cells were collected by trypsinization, washed in PBS, fixed in 70% ice-cold ethanol, rehydrated in PBS and stained with Propidium Iodide (40μg/ml) for 30 min at room temperature. The DNA content of cells stained by propidium iodide was analyzed by a BD LSR II flow cytometer (BD Biosciences). For analysis of mitotic cells by pH3-Ser10 staining cells were collected similarly, followed by permeabilization with 0.25% Triton for 15 min, washed with 1% BSA and incubated with α-pH3-Ser10 in 1% BSA for 2h and with Donkey-α-Rabbit-FITC F(ab’)2 in 1% BSA for 30 min. Cells were analyzed by a BD LSR II Flow Cytometer. For microscopic flow cytometry analysis cells were fixed in 1% PFA (paraformaldehyde) in PBS (phosphate-buffered saline), immunostained and analyzed using an ImageStream Imaging Flow Cytometer.

### Immunoblotting

Whole cell lysates were prepared by incubating PBS-washed cells in lysis buffer containing 1% NP-40, 1% sodium deoxycholate, 0.1% SDS, 150 mM NaCl, 50 mM Tris-HCl pH 7.4, 1 mM EDTA, 1 mM PMSF (RIPA buffer) and a protease inhibitor cocktail (Sigma) and phosphatase inhibitor cocktails II and III (Sigma, for Tyrosine and Serine/Threonine phosphatases). Lysis was performed by incubation of cells for 10 min on ice followed by sonication for 30s at intervals of 10s at 4 °C. Lysates were then centrifuged at 13 000*g* for 15 min followed by discarding of the insoluble pellet. Protein concentration was determined using the Bradford protein assay (Bio-Rad). Following addition of β-mercaptoethanol-containing sample buffer to equal amounts of protein, samples were boiled for 3 minutes and resolved by SDS-PAGE (8% acrylamide) and electrophoretically transferred onto polyvinylidene difluoride membranes (Immobilon-P, Millipore, Bedford, MA). For detection of phosphorylated histones, whole cell lysates were prepared by incubation in RIPA buffer at 4°C for 20 minutes, followed by sonication and separation by SDS-PAGE (15% acrylamide). Transferred membranes were blocked with Tris-buffered saline containing 0.5% nonfat dry milk and 0.1% Tween 20 for 1hr at room temperature. After blocking, the membranes were incubated with the appropriate primary antibodies for 1 hr room temperature or at 4°C overnight. Membranes were then incubated with the appropriate secondary antibodies for 1 h at room temperature. Immunoblots were visualized by enhanced chemiluminescence, using the EZ-ECL detection reagent (Biological Industries, Kibbutz Beit Haemek, Israel) according to the manufacturer’s instructions, and signals were detected by the ImageQuant LAS 4000 (GE Healthcare) or by exposure to film.

### Immunoprecipitation

For immunoprecipitation of myc-Plk1, cell extracts were prepared by lysis of PBS-washed cells in RIPA buffer (1% NP-40, 1% sodium deoxycholate, 0.1% SDS, 150 mM NaCl, 50 mM Tris-HCl pH 7.4, 1 mM EDTA, 1 mM PMSF) at 4°C for 10 minutes. The insoluble pellet was discarded following centrifugation for 15 min at maximal speed and protein concentration was determined using the Bradford protein assay (Bio-Rad). Samples were incubated with an α-myc antibody overnight followed by incubation with protein A/G agarose beads for 2hr and washes. Following addition of β-mercaptoethanol-containing sample buffer samples were boiled for 5 minutes and the precipitated proteins were separated by SDS-PAGE (8% acrylamide) followed by immunoblotting with the appropriate antibodies.

### G2/M checkpoint release assay

Cells were seeded at a density of 800,000 cells per 6 cm plate. Thymidine was added, later the same day, at a concentration of 2.5mM. Following a 24hr incubation, thymidine was washed away, and cells were irradiated or treated with Doxorubicin 6 hours later (followed by wash of Doxorubicin after 1.5h). Nocodazole was then added for 16h (overnight). On the next morning, caffeine (5mM) was added for 9-12 hrs, until cells began detaching from the plate. Cells were then harvested for both immunoblotting and FACS analysis, followed by staining with α-pH3-Ser10 to detect mitotic cells.

### shRNA mediated knockdown

For stable knockdown of c-Abl, HEK293T cells were transfected, using the calcium phosphate method, with mission pLKO.1 shRNA vectors targeting c-Abl or GFP (Sigma) together with pLP1, pLP2, and pLP-VSVG (vesicular stomatitis virus glycoprotein) packaging plasmids (Invitrogen). Forty-eight hours after transfection, viral supernatant was removed and filtered through a 0.45-μm filter, supplemented with 8 μg/mL polybrene, and used to infect U2OS cells. Twenty-four hours after infection, infected cells were selected with the addition of 2 μg/mL puromycin (Sigma) to the culture medium.

### Staining & Microscopy

Nuclear staining of live cells was performed with Hoechst-H33342 for 15 min at 37 °C. For analysis of nuclei of fixed cells, cells were seeded on glass coverslips. Fixation was performed in 4% paraformaldehyde for 30 minutes at room temperature. Coverslips were mounted using DAPI Fluoromount-G (Southern Biotech, AL, USA). Slides were analyzed with a Zeiss LSM 710 Meta confocal microscope using ZEN software.

### Quantification of Micronuclei formation

Cells were grown on glass coverslips and treated with IR to induce micronuclei formation, alongside unirradiated control samples. Cells were fixed with 4% paraformaldehyde 24 hours post-IR. Each sample was collected in duplicate to ensure data reproducibility. Images were acquired using a Zeiss LSM 710 Meta confocal microscope. Micronuclei were counted using a MATLAB-based program (MNiDcount) designed to facilitate the counting process. The program enables efficient navigation between captured images, identifies cell nuclei, and provides an interface for user-supervised classification, annotation, and quantification of micronuclei and other post-mitotic nuclear abnormalities. For each sample, at least 1,000 nuclei were analyzed.

### In Vitro Kinase Assay

To assess the activity of Plk1 kinase, HEK-293 cells were transfected with myc-Plk1 wild-type (WT) or myc-Plk1 Y217F or myc-Plk1 K82R (kinase-dead Plk1) constructs, together with an active form of c-Abl kinase (c-Abl Δ1-81) or its corresponding kinase mutant (c-Abl Δ1-81 km), as indicated. Cells were harvested 24 hours post-transfection and lysed in RIPA buffer. Immunoprecipitation of Plk1 was performed as described using an α-myc antibody. The immunoprecipitates were incubated in a kinase reaction buffer (50 mM Tris-HCl, pH 7.5, 10 mM MgCl2, 1 mM EGTA, 2 mM DTT, and 0.01% Brij 35) containing purified Casein kinase (CSK) as a substrate, and (γ-^32^P)-ATP (PerkinElmer, Waltham, MA, USA). Reactions were incubated at 37 °C for 60 min and terminated by the addition of SDS sample buffer, followed by boiling. Samples were resolved by SDS-PAGE and transferred onto PVDF membranes (Millipore Corp., Billerica, MA.) To evaluate ^32^P incorporation, the membrane was dried and exposed to a PhosphorImager screen for 3 hours, then analyzed using a PhosphorImager system (Amersham Biosciences). To confirm that the observed differences in the radioactive signal were not due to variations in protein levels, protein levels were assessed by immunoblotting for Plk1 expression (IB: α-myc)

### CRISPR-mediated genome editing

Two guide RNAs were designed against Plk1 Exon3 (gRNA1 – 5’TCCTAATTACATAGCTCCCG3’, gRNA2 – 5’AGAGGAAGAAGACCCTGTGT3’) and cloned into a pX330 backbone. An ssODN of 122 nucleotides homologous to a sequence within Plk1 Exon3 was designed to contain either of the following mutations - Y217F, Y217E, T210A, T210D - as well as mutations within the NGGs of the guide RNA target sequences. ssODNs Y217F and Y217E were transfected into U2OS cells together with guideRNA1. ssODNs T210A and T210D were transfected into U2OS cells together with guideRNA2. An expression plasmid for GFP was added to the transfection. 24 hours following the transfection GFP positive cells were sorted by FACSort cytometer under sterile conditions and seeded on small plates to enlarge stock. Cells were then grown under normal conditions. Presence of the inserted Plk1 cassette was confirmed by PCR with mutation-specific primers.

Some sections of the experimental outlines were created using BioRender.com

## Acknowledgments

We sincerely thank Alexander Meltzer for designing and developing the MNiDcount program. His expertise and dedication in creating a user-friendly and efficient tool for micronuclei quantification have greatly enhanced our data analysis process. We thank Dr. Michael Brandeis for the myc-Plk1 vector. Research was supported by a grant from the Estate of Gertrude Buchler and the Estate of Hermine Miller.

## Supplemental Information

To assess the role of c-Abl kinase activity in facilitating G2/M transition following IR-induced DNA damage, HEK293 cells were transfected either with a constitutively active form of c-Abl (c-Abl Δ1-81), which lacks the regulatory N-terminal domain, or a kinase-dead c-Abl mutant (c-Abl Δ1-81 Km) (12). After transfection, cells were irradiated, and cell cycle progression was analyzed 24 hours post-IR. Notably, cells transfected with either an empty vector or the kinase-dead c-Abl mutant exhibited significant G2/M accumulation. In contrast, cells transfected with constitutively active c-Abl showed no such accumulation (Fig. S1A), strongly suggesting that c-Abl kinase activity is associated with reduced G2/M arrest following DNA damage.

To validate that the prolonged G/M accumulation upon Imatinib treatment is c-Abl dependent, cell cycle analysis was performed following IR-induced DD damage in wild-type (WT) MEFs and c-Abl null (c-Abl-/-) MEFs. While Imatinib treatment led to G2/M accumulation in WT MEFs following IR, c-Abl-/- MEFs showed no significant G2/M accumulation under the same conditions (Fig. S1B), confirming that the Imatinib effect on DD-induced G2-M arrest is c-Abl dependent.

To determine whether c-Abl actively contributes to G/M checkpoint release, we employed shRNA-mediated c-Abl knockdown in U2OS cells and CRISPR-mediated c-Abl knockout in HEK293FT cells. Depletion of c-Abl across three distinct CRISPR knockout clones, was confirmed by immunoblot (Fig. S1C). Similarly, c-Abl knockdown using two distinct shRNAs, each targeting different regions of c-Abl mRNA, was validated by immunoblotting (Fig. S1D).

To determine whether c-Abl regulates release from G2/M arrest, we applied a G2/M checkpoint release assay (Fig. S2A), adapted from (36). Cells were synchronized with thymidine for 24 hours, followed by thymidine wash-off and subsequent DNA damage induction by IR or doxorubicin. One hour after DNA damage, microtubule polymerization inhibitor nocodazole was added to prevent mitotic progression, and after 15-hours, the cells were arrested in G2/M (Fig. S2B, small histogram insets). To facilitate synchronous release from G2/M arrest, cells were treated with caffeine and analyzed for mitotic entry by assessing the proportion of 4N cells positive for the phospho-histone H3 Ser10 mitosis marker (37). Following nocodazole treatment, nearly half of the unstressed cycling cell population exhibited strong pH3-(Ser10) staining, indicating mitotic entry followed by entrapment in mitosis. In contrast, IR-treated cells displayed minimal mitotic entry (Fig. S2B). Upon caffeine treatment, irradiated cells showed a gradual increase in pH3-(Ser10) staining, with significant mitotic entry observed 6 hours post-treatment with caffeine (Fig. S2B). Immunoblot analysis confirmed mitotic entry commencing approximately 5-6 hours after caffeine addition, as evidenced by increased histone H3 Ser10 phosphorylation and decreased Chk1 Ser345 phosphorylation, both markers of G2/M checkpoint release (Fig. S2C). Caffeine treatment also resulted in reduced levels of Claspin and Wee1, proteins essential for G2/M checkpoint induction and maintenance, whose degradation is crucial for G2/M checkpoint release (38) (Fig. S2D). To assess cell cycle distribution prior to caffeine addition, identical cell samples from the checkpoint release assay in Fig. 2B and Fig. 2C, but without Nocodazole treatment, were analyzed (Fig. S2E and 2F).

To assess the kinetics of Plk1 tyrosine phosphorylation following IR-induced DD, HEK293 cells were transfected with myc-Plk1 WT and irradiated 24 hours post-transfection. Cells were harvested at multiple time points post-IR and lysates were subjected to immunoprecipitation with α-myc, followed by immunoblotting with the phosphotyrosine-specific antibody PY20 (Fig. S3A). A notable increase in Plk1 tyrosine phosphorylation was detected beginning around 12 hours and peaking at 18-20 hours post-IR, coinciding with the onset of G2/M checkpoint release and mitotic entry. These findings suggest that Plk1 tyrosine phosphorylation may be functionally important for G2/M checkpoint release.

To determine whether c-Abl directly phosphorylates Plk1, myc-Plk1 WT was co-expressed with increasing amounts of constitutively active c-Abl (c-Abl Δ1-81). Immunoprecipitation with α-myc followed by PY20 immunoblotting revealed dose-dependent phosphorylation of Plk1 by c-Abl (Fig. S3B).

**Figure S1.**
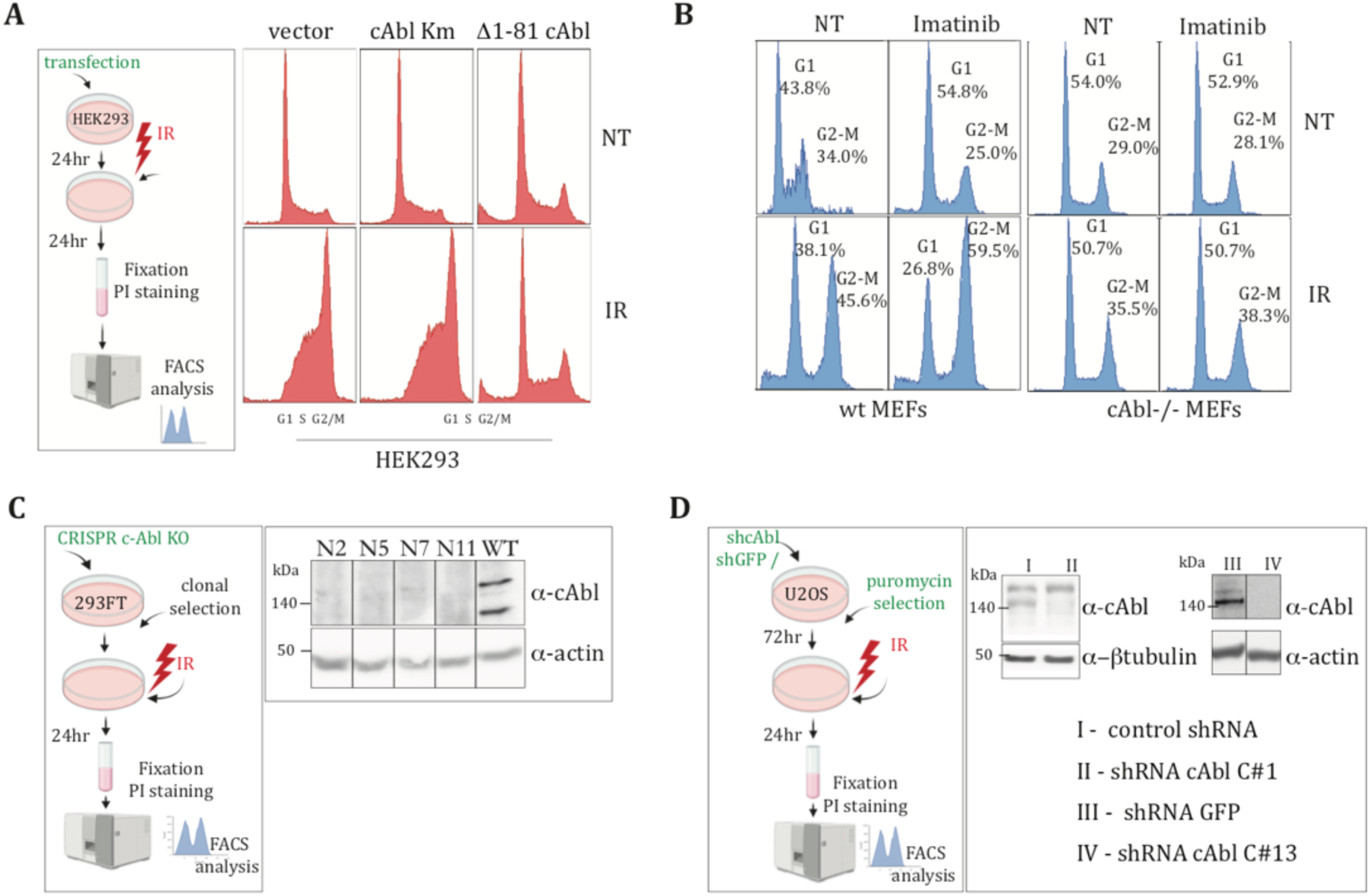
**(A)** HEK293 cells were transiently transfected with either an empty vector or constructs expressing a constitutively active c-Abl kinase (Δ1-81 c-Abl) or kinase-defective c-Abl mutant (c-Abl Km). Following irradiation, cell cycle distribution was analyzed by FACS 24 hours post-IR. **(B)** Wild-type (WT) and c-Abl-null (c-Abl-/-) MEFs were pretreated with Imatinib for 2hours before IR exposure. Cells were then fixed, stained with propidium iodide (PI) and analyzed by FACS 24 hours post-IR to assess cell cycle distribution. **(C)** CRISPR-mediated knockout of c-Abl was performed in HEK293 cells. Four independent clonal lines were validated by immunoblotting to confirm knockout efficiency **(D)** shRNA-mediated knockdown of c-Abl in U2OS cells was carried out using a third-generation lentiviral system. Cells were harvested 72 hours following infection with shRNA-expressing lentivirus and puromycin selection. c-Abl knockdown efficiency was confirmed by immunoblot in two distinct shRNA clones.

**Figure S2.**
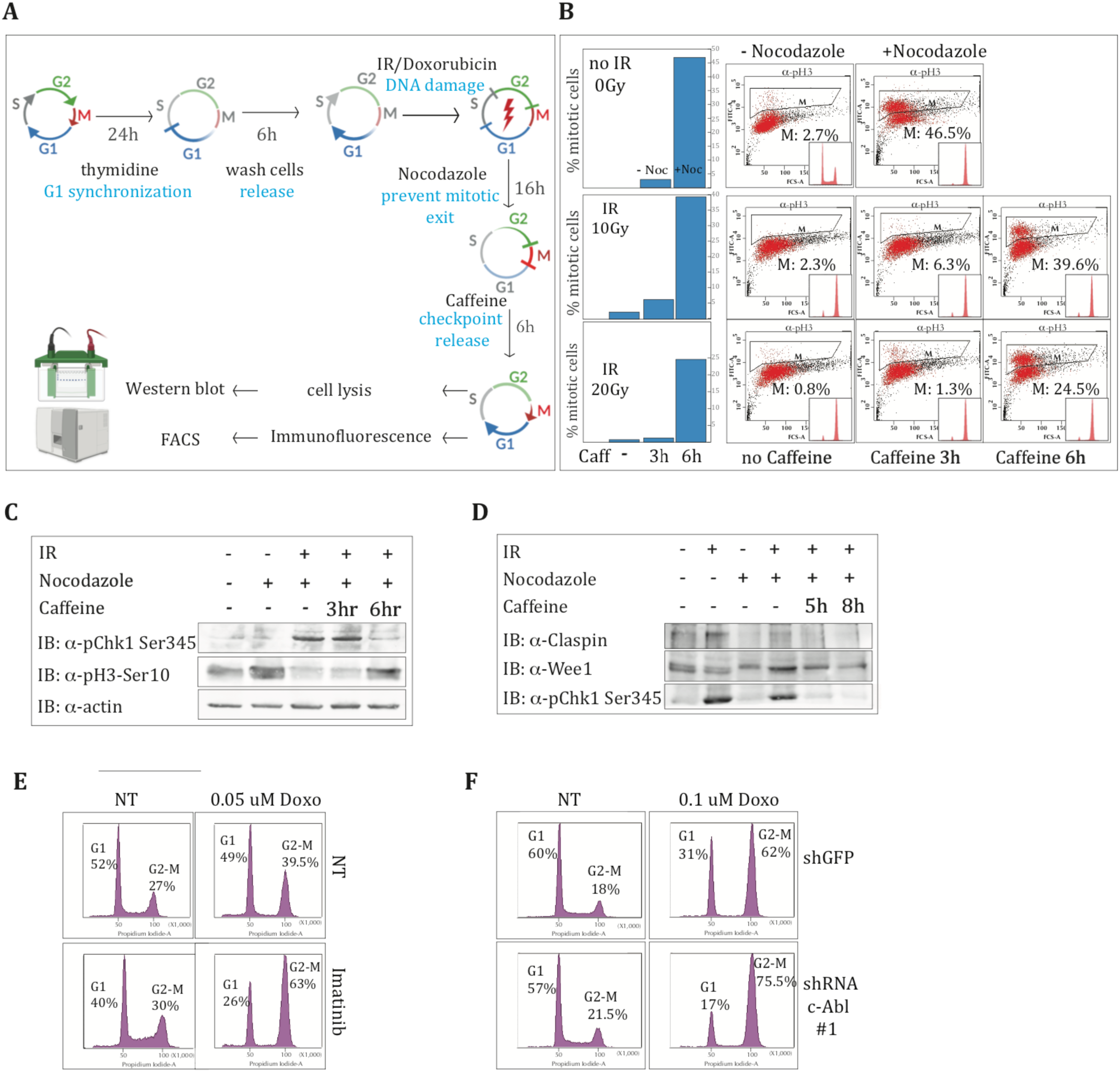
**(A) G2/M checkpoint release assay - experimental setting.** Cells were synchronized in G1 by a 24-hour incubation with thymidine, then released to enter S phase. Following 6 hours of release from thymidine arrest, DNA damage was induced either by ionizing radiation (IR) or by incubation with doxorubicin for 1 hour, followed by medium exchange to remove the drug. Next, nocodazole was added to prevent mitotic exit of cells that have completed G2. Following 16 hours, cells were treated with caffeine to override G2-M checkpoint maintenance by inhibition of the ATM/ATR kinases and allow assessment of the effectiveness of G2/M checkpoint release. Cells were collected 6-12 hours following caffeine addition for immunoblot and phospho-histone H3 (pH3-Ser10) flow cytometry analysis **(B) Validation of the G2/M checkpoint release assay.** Representative FACS plots of α-pH3 (Ser10) staining following the indicated treatments. The columns of the left show quantification of the pH3 (Ser10) signal (by the FACSDiva software). The top row shows validation of mitotic entry in the absence of DNA damage, following nocodazole treatment, while the middle and bottom rows illustrate checkpoint release kinetics following addition at IR doses of 10 Gy and 20 Gy, respectively. **(C)** Immunoblot analysis with α-p-Chk1 Ser345 and α-phospho-H3 following the checkpoint release assay. **(D)** Immunoblot with α-Claspin and α-Wee1 following the checkpoint release assay. **(E)** Thymidine-synchronized HeLa cells were pretreated with Imatinib or left untreated, followed by treatment with doxorubicin for 1 hour. Cell cycle distribution was subsequently analyzed by FACS. **(F)** Thymidine-synchronized c-Abl knockdown (shcAbl #1) and control knockdown (shGFP) U2OS cells were treated with doxorubicin for 1 hour. Cell cycle distribution was analyzed by FACS.

**Figure S3.**
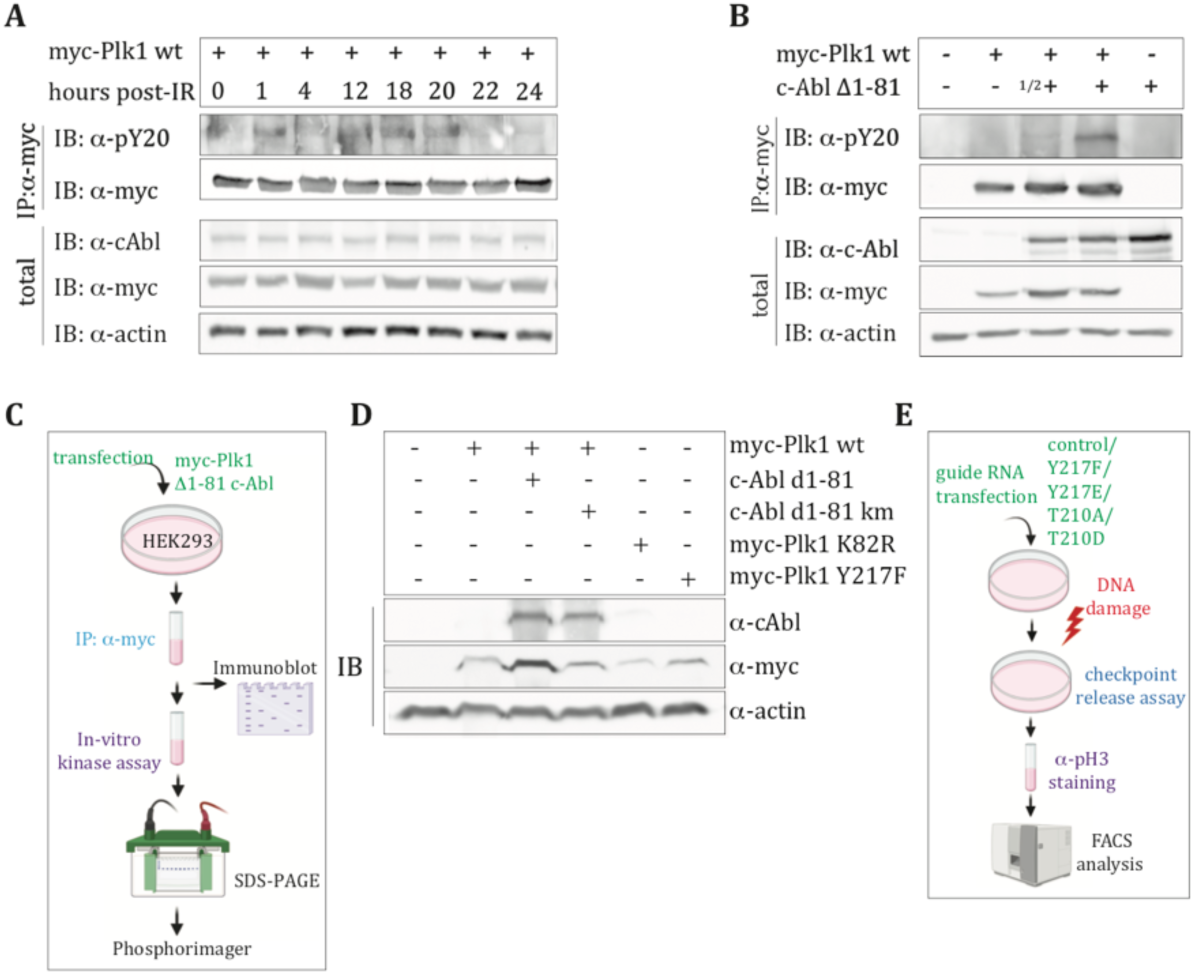
**(A)** HEK293 cells transfected with myc-Plk1 WT were irradiated (10Gy) 24 hours post-transfection and collected at the indicated time points. Plk1 was immunoprecipitated using α-myc and immunoblot analysis was performed with the indicated antibodies. An α-phosphotyrosine (TY20) antibody was used to detects overall tyrosine phosphorylation and monitor Plk1 tyrosine phosphorylation kinetics following IR-induced DNA damage. **(B)** HEK293 cells were transfected with myc-Plk1 and with increasing amounts of constitutively active c-Abl (c-Abl Δ1-81). At 24 hours post-transfection, Plk1 was immunoprecipitated (α-myc) and analyzed by immunoblotting with the indicated antibodies to assess Plk1 tyrosine phosphorylation in presence of active c-Abl kinase**. (C)** Experimental outline of the in-vitro kinase assay. **(D)** Expression levels of the constructs used in the in-vitro kinase assay (Fig.2F) were confirmed by immunoblotting (α-myc) in parallel samples. All constructs were transfected at equivalent amounts. **(E)** Schematic overview of the CRISPR-mediated knock-in strategy for introducing Plk1 mutations

**Figure S4.**
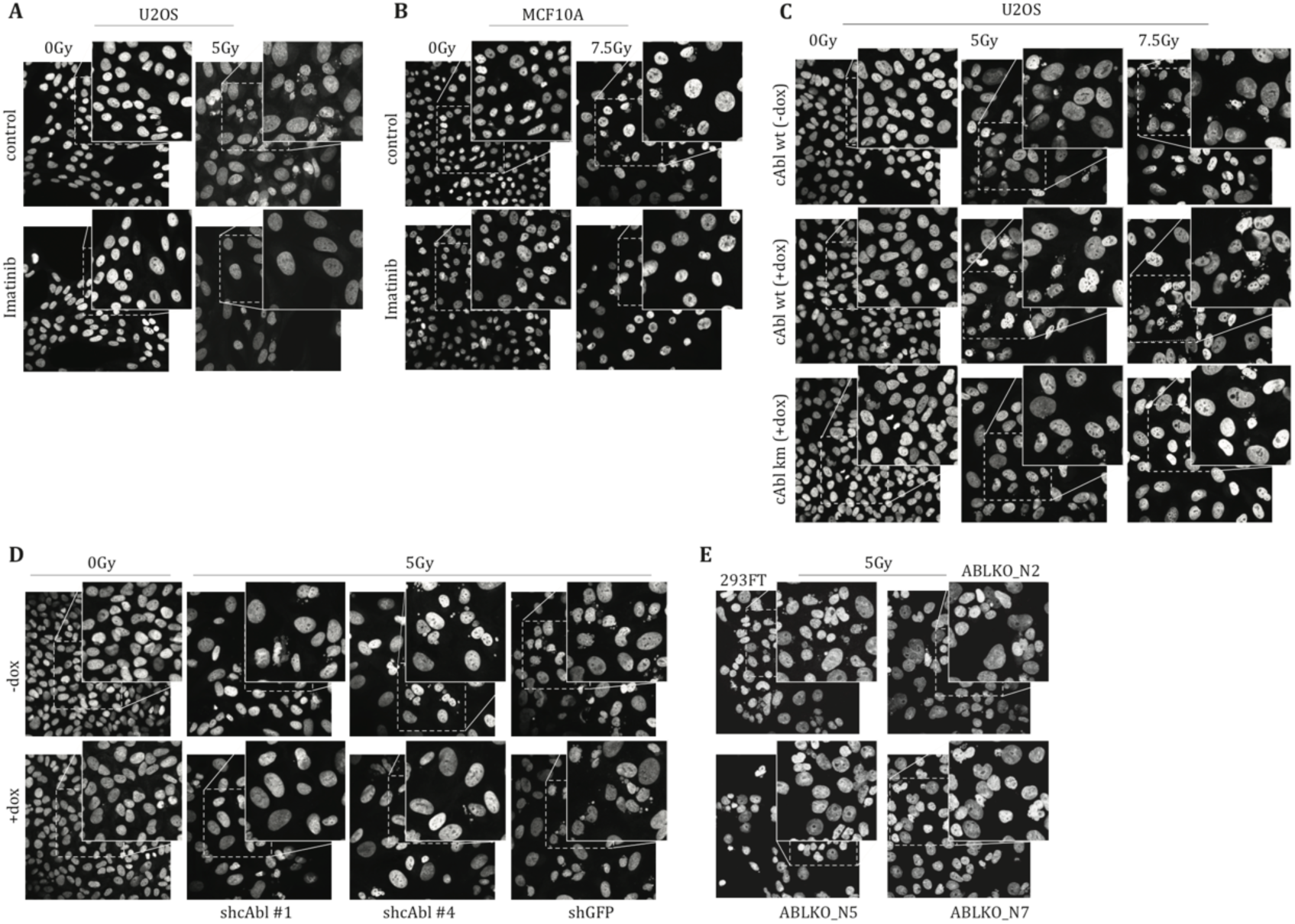
**(A)** Representative fluorescence images of U2OS cells from experiment shown in Fig.5A **(B)** Representative fluorescence images of MCF10A cells from the experiment shown in Fig.5B **(C)** Representative images of doxycycline-inducible U2OS cells from the experiment shown in Fig.5C **(D)** Representative images of doxycycline-inducible shRNA U2OS cells from the experiment shown in Fig.5D. **(E)** Representative images of CRISPR-generated HEK293FT c-Abl knockout clones from the experiment shown in Fig.5E.

## Notes

### Competing Interest Statement

The authors have declared no competing interest.

